# Impact of an alpha helix and a cysteine-cysteine disulfide bond on the resistance of bacterial adhesion pili to stress

**DOI:** 10.1101/2021.01.18.427124

**Authors:** Joseph L. Baker, Tobias Dahlberg, Esther Bullitt, Magnus Andersson

## Abstract

*Escherichia coli* express adhesion pili that mediate attachment to host cell surfaces that are exposed to body fluids in the urinary and gastrointestinal tracts. Pilin subunits are organized into helical polymers, with a tip adhesin for specific host binding. Pili can elastically unwind when exposed to fluid flow force, reducing the adhesin load, thereby facilitating sustained attachment. Here we investigate biophysical and structural differences of pili commonly expressed on bacteria that inhabit the urinary and intestinal tracts. Optical tweezers measurements reveal that Class 1 pili of uropathogenic *E. coli* (UPEC), as well as Class 1b of enterotoxigenic *E. coli* (ETEC), undergo an additional conformational change beyond pilus unwinding, providing significantly more elasticity to their structure than ETEC Class 5 pili. Looking comprehensively at structural and steered molecular dynamics simulation data, we find this difference in Class 1 pili subunit behavior originates from an *α*-helical motif that can unfold when exposed to force. A disulfide bond cross-linking *β*-strands in Class 1 pili stabilizes subunits, allowing them to tolerate higher forces than Class 5 pili that lack this covalent bond. We suggest that these extra contributions to pilus resiliency are relevant for the UPEC niche since resident bacteria are exposed to stronger, more transient shear forces compared to those experienced by ETEC bacteria in the mucosa of the intestinal tract. Interestingly, Class 1b ETEC pili include the same structural features seen in UPEC pili, while requiring lower unwinding forces that are more similar to those of Class 5 ETEC pili.

**Significance Statement:** Adhesion pili are often essential virulence factors for attachment of pathogenic bacteria in specific environmental niches. We provide mechanistic details of structural differences impacting the biophysical properties of pili found on bacteria in the urinary and intestinal tracts. We see that pili from urinary tract bacteria are composed of subunits optimized for their microenvironment. First, they can tolerate higher forces than intestinal pili due to a disulfide bond that limits subunit unfolding. Second, their greater flexibility is due to an *α*-helical motif that can unfold, absorbing force that could otherwise lead to bacteria detachment. Our work provides insight into the central role of pilus structural and biophysical properties for the sustained bacterial adherence necessary to initiate disease.

---

*Escherichia coli (E. coli)* have a remarkable ability to adapt to the environment, allowing these bacteria to colonize varying niches in humans and animals either as commensals or pathogens (1). The urinary and gastrointestinal tracts are examples of environments where pathogenic *E. coli* are common causes of acute urinary tract infections and severe diarrhea, respectively. In these niches, fluid flow is a natural defense mechanism, limiting attachment of pathogenic bacteria to epithelial cell surfaces (2, 3). To facilitate attachment under fluid flow, *E. coli* use attachment organelles called adhesion pili or fimbriae (4). Adhesion pili of uropathogenic *E. coli* (UPEC) and enterotoxigenic *E. coli* (ETEC) pili that are assembled via chaperone-usher pathways are micrometer long helical rod structures, with an adhesin at the tip that binds to host receptors (Figure 1A) (5). The helical rod structure can be unwound, significantly extending its original length under tensile force. This unwinding allows cell-associated bacteria to withstand shear forces from fluid flow, by decreasing the load on the receptor-bound adhesin (6, 7). This unwinding of pili is dependent on critical mechanical features of the fibers. If the pili are compromised, the bacteria’s ability to attach and stay attached under shear force is reduced significantly (8, 9), and ETEC with no pili are unable to cause disease (10). UPEC and ETEC pili mechanics and structure have been investigated for decades, yet we still lack a complete picture of their mechanical differences and how these differences relate to their genetics, structure and environmental niche.

**Fig. 1.**
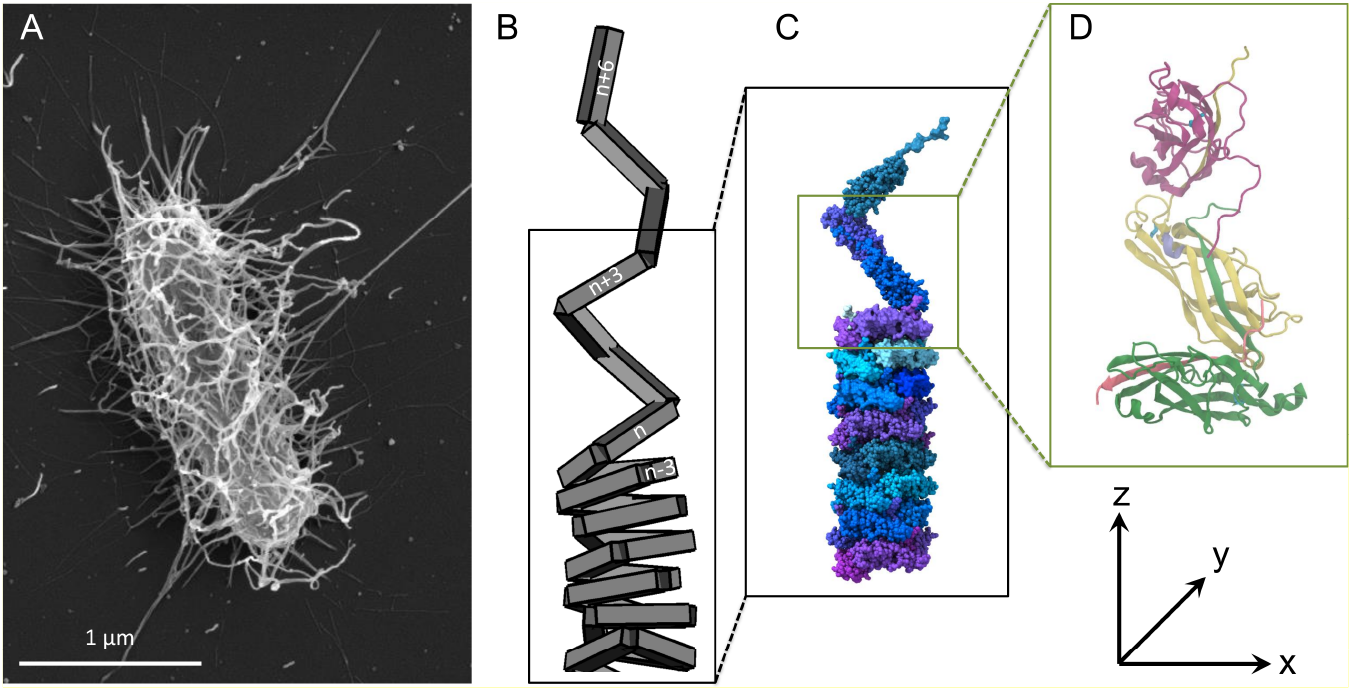
A) Scanning electron microscopy image of an *E. coli* bacterial cell expressing P pili. B) Cartoon of a pilus showing subunits assembled into a helix-like rod stabilized by layer-to-layer interactions between subunits n and n±3, and a region of unwinding. C) High-resolution structure of a P pilus with each pilin subunit colored individually. D) a trimer (3mer) of pilin subunits during unwinding.

UPEC and ETEC pilus rods and pilin subunits are structurally similar, while there are genetically distinct classes of pilins (Class 1 and Class 5). Structurally, both pilus types are composed of immunoglobulin (Ig)-like pilin subunits attached via a *β*-strand complementation head-to-tail, forming helical fibers of approximately 1,000 pilins with similar quaternary structure (11, 12). The stability of the quaternary helical rod structure is achieved via layer-to-layer bonds formed between subunits, primarily between subunits *n* and *n* + 3 (Figure 1B). High-resolution three-dimensional helical reconstruction models exist for UPEC Class 1 Type 1 and P pili (Figure 1C-D), whereas only CFA/I pili of ETEC has been reconstructed at a resolution sufficient for atomic model building (9, 13–15). These reconstructions show that UPEC pili have a larger buried surface area between subunits *n* and *n* + 3 than ETEC pili (1616 Å^2^ and 1453 Å^2^ *vs.* 1087 Å^2^, respectively). The magnitude of the buried surface area correlates well with the force needed to unwind the fibers. That is, larger buried surface area requires a higher tensile force: the force needed to unwind Type 1 and P pili is more than fourfold that of CFA/I pili, ≈ 30 pN and 28 pN *vs.* 7 pN (15, 16). Overall, the unwinding capabilities of pili are well understood, and good biophysical models explain the measured force-extension curves (17–20). However, current force-extension data indicate that there is a puzzling difference in the mechanics of pili that these models cannot explain, that could be related to pilin sequence differences. Class 1 pili show an additional conformational change that takes place at almost twice the unwinding force (≈ 60 pN), which makes them more elastic and allows significantly longer extensions than those of Class 5 pili that were studied previously (16, 21–23). Since the conformational change in Class 1 pili takes place after unwinding of the quaternary structure, it must occur when a pilus is already in its linearized form (subunits in a head-to-tail order). That is, after unwinding the helical rod, there are changes in secondary or tertiary structures of individual pilin subunits. However, these changes have not been explored.

It is well established that UPEC pilin subunits are proteins with high mechanical stability. Pilin stability originates from the Ig-like structure that is assembled of six *β*-strands forming a *β*-sandwich, and includes a conserved disulfide bond that work as a mechanical lock (24, 25). These physical attributes yield pilins that are characterized by very high thermodynamic and kinetic stability; free energies of over 70 kJ/mol and a half-life of 10^8^ years at 25x°C (24, 26), as well as being robust under tensile force (25). In contrast, little is known regarding the physical attributes of ETEC pilin stability, except that there are striking similarities with UPEC pilins regarding their IgG protein fold and size (23). Thus, we raise the following question: what provides the mechanical differences observed between UPEC and ETEC adhesion pili, and is this difference universal?

To solve the aforementioned research question, we compared the mechanical differences and structural properties of UPEC- and ETEC-associated pili. We measured their mechanical properties using optical tweezers force spectroscopy and we interpret the experimental force-extension results in the final region of pilus extension using structural models and steered molecular dynamics (sMD) simulations. We also look closely at Class 1 ETEC pili that have more genetic homology to UPEC pili as compared to Class 5 ETEC, to examine the relation between genetics, pilus mechanics, and environmental niche.

## 1. Results

### A. Optical tweezers measurements reveal a significant difference in Class 1 vs Class 5 pili elasticity

To investigate the biophysical and structural differences between polymeric pili expressed in each respective host niche, we used optical tweezers to measure their mechanical properties. A measurement was performed by first trapping a bacterial cell with the optical tweezers and attaching the cell to a 9.5 *μ*m diameter poly-l-lysine functionalized microsphere that was immobilized on a glass slide (27). We subsequently trapped a two *μ*m diameter polystyrene microsphere and non-specifically attached it gently to the end of a pilus. We extended the pilus by applying a tensile force, keeping the trap fixed and moving the glass slide with the immobilized microsphere at a constant speed that fulfilled the criterion of steady-state conditions of adhesion pili (19); steady-state extension permits assessment of the unwinding force without the contribution of a dynamic force response. Finally, we point out that this study presents the first steady-state response of Type 1 pili, since previous measurements on these structures all were assessed in a dynamic regime (6, 16, 28). Details of the instrumentation and measurement procedure are found in the Methods section and Supporting Information.

Using the same procedure we examined three pilus types from the urinary tract and three from the intestinal tract niches: P, Type 1, and F1C pili; and CFA/I, CS2, and CS20 pili expressed on bacteria that cause UPEC or ETEC infections, respectively. As seen for all representative force-extension data curves in Figure 2 panels (A-F), all pili showed a force that first linearly increased in region I, followed by an approximately flat force plateau in region II, whereafter it again increased in region III. For UPEC, panels (A-C), this increase in region III was a combination of three linear regions with smooth transitions; that is, the shape is sigmoidal. The sigmoidal shape, in which the central region rises more slowly rather than continuing linearly, is an indication of stochastic conformational changes (17, 29). Also, note that for panels (A-C) this transition starts at approximately 50 pN, which is 60 % higher than the unwinding force. Conversely, for two ETEC pilus types, panels (D-E), the increase in region III is purely linear. We then measured the force-extension for CS20 pili, an ETEC pilus that is comprised of a pilin that is genetically similar to UPEC, and is therefore a Class 1 pilus. As seen in Figure 2F, our data showed the sigmoidal increase seen in its Class 1 homologues, rather than having the purely linear rise seen in the other ETEC pili, which are comprised of Class 5 pilins. That is, UPEC and ETEC pili showed similar responses for regions I and II, and when further stretched (region III), both types of pili initially increased linearly in force. With continued extension, all UPEC pili and ETEC CS20 pili showed a transition in the force-extension curve related to a stochastic conformational change, as seen by the more slowly increasing force plateau. This elastic change significantly increased the pili’s length by 25 % compared to a strictly linear region III force-extension curve. Since this transition took place when the pilus filament was already in a linearized form, we speculated that this conformational change occurred either within individual pilin subunits or in the head-to-tail bond between pilins. To identify and clarify this research question we turned to structural analysis.

**Fig. 2.**
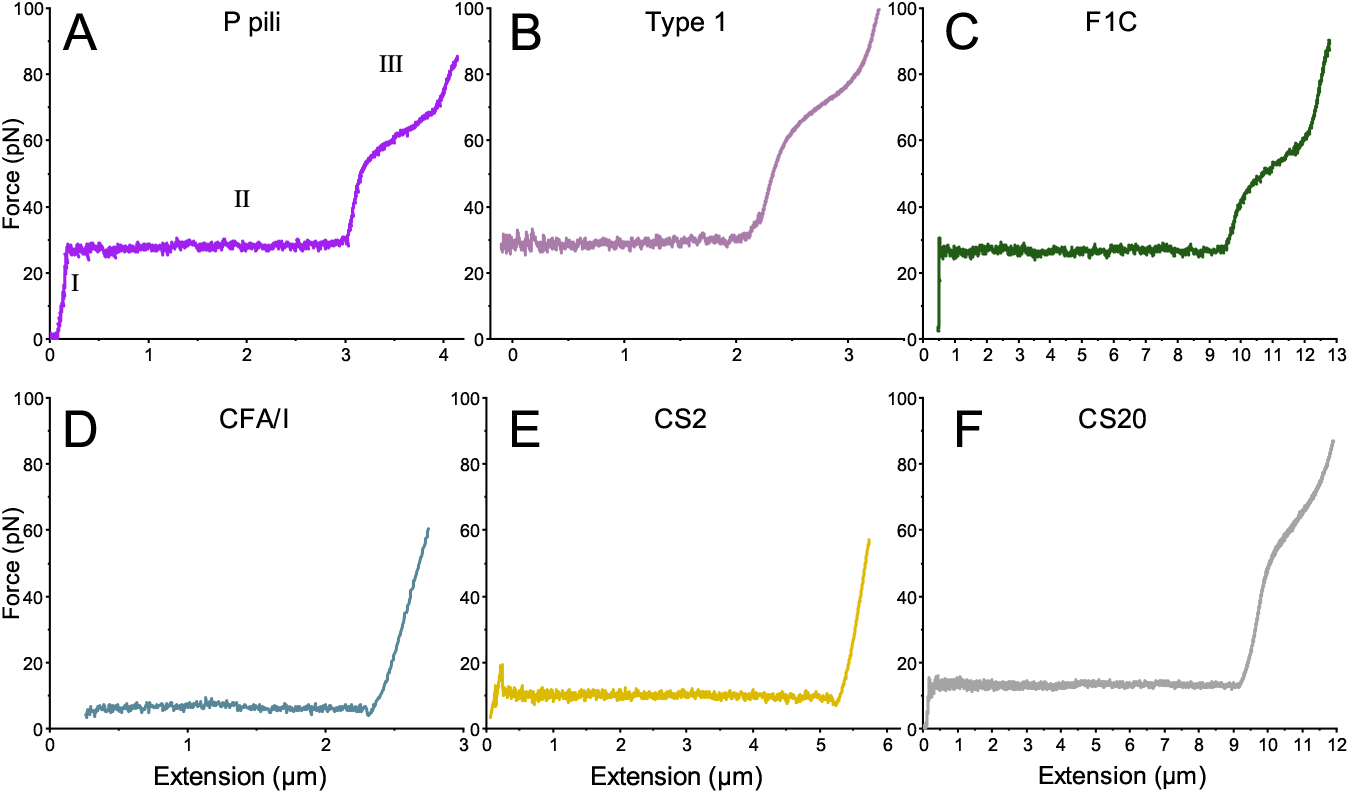
Optical tweezers force-extension curves of pili commonly found on bacteria in the urinary tract (upper row) and the intestinal tract (lower row). The three different regions are defined in panel A and we give for each pilus type the region II unwinding force (population mean±standard deviation) A) P pilus 28±1 pN (n=30) B) Type 1 pilus 30±1 pN (n=30) C) F1C pilus 26±2 pN (n=30) D) CFA/I pilus 7±1 pN (n=30) E) CS2 pilus 10±1 pN (n=30) F) CS20 pilus 15±1 pN (n=30). These unwinding forces correspond well with previously published values (16, 22, 23, 30, 31). Note that since pili were attached via non-specific interactions and the different pili examined have different hydrophobic surfaces, pili detached from the microsphere at different forces. We observed that in general, the ETEC-expressed pili were more difficult to attach to microspheres and that they detached at a lower force. This however, does not affect the shape of the force response, only the amount of applied force prior to detachment.

### B. Structural analysis reveals an *α*-helical motif and a stabilizing disulfide bond in Class 1 pili not present in Class 5 pili

We examined the pilins’ structures to interpret the source(s) of difference in optical tweezers force-extension measurements in region III force curves of UPEC *vs* ETEC, and Class 1 *vs* Class 5 pili, as shown in Figure 3. The experimentally determined UPEC Class 1 pilin structures, PapA and FimA, and the homology-modeled FocA, all showed the presence of a small *α*-helical motif in the structures of their pilins. The location of the *α*-helix was approximately 20 amino acids from the N-terminus. For example, in PapA (pdb: 5FLU) this *α*-helix includes residues 26-29 (see Supporting Information Table S1 for all Class 1 *α*-helix residues). With respect to Class 5 ETEC pili, an open turn was visible in the structures in this region, but no *α*-helical secondary structure was present in the known structure of the CFA/I pilin CfaB (pdb: 6NRV), nor in the homology-modeled Class 5 pilin CotA (Figure 3) as determined using the DSSP structure determination algorithm in ChimeraX (32).

**Fig. 3.**
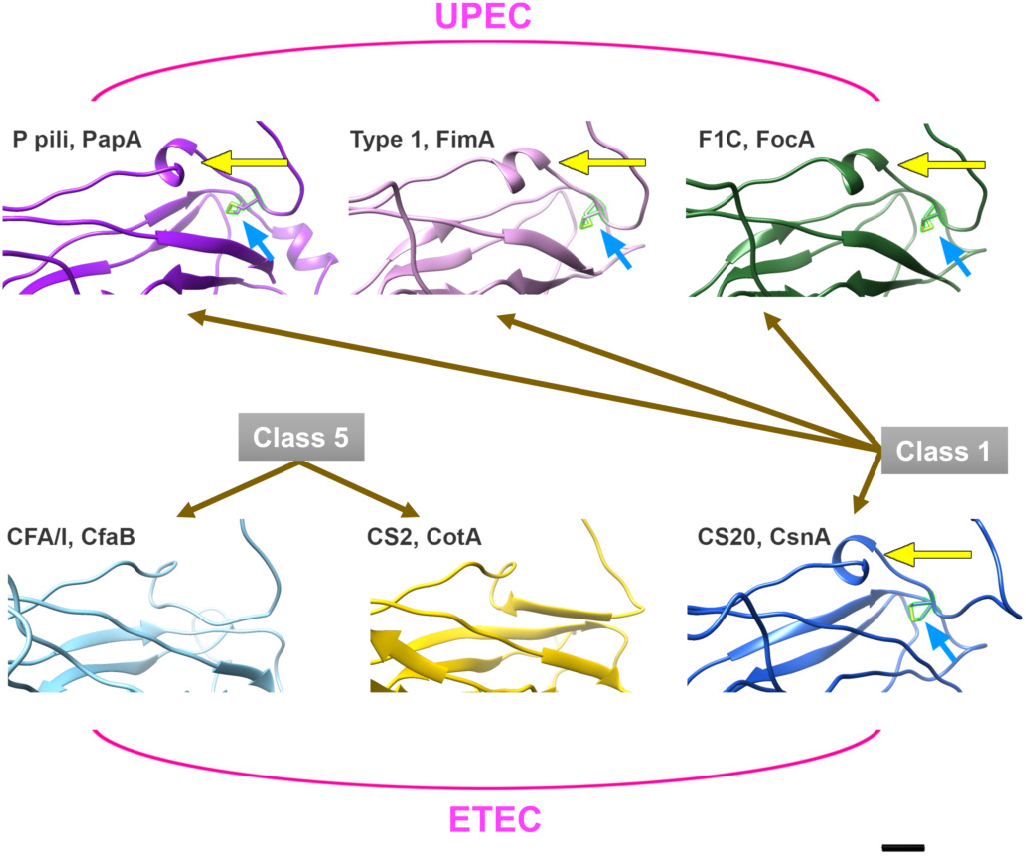
Structural analysis of pilin subunits. Top row: Class 1 pilins from UPEC (PapA, FimA, and FocA) all include a short *α*-helix near the N-terminus of each pilin (yellow arrows). In addition, a disulfide bond between two cysteine residues links together two *β*-strands within the subunit (blue arrows, and S-S highlighted in green). Bottom row: Class 5 pilins from ETEC (CfaB and CotA) have an open loop near the N-terminus, but no short *α*-helical structures are present. Class 5 pilins have no cysteines, and thus there are no disulfide bond interactions. Class 1 CsnA pilins from ETEC CS20 pili have the same structural features as UPEC Class 1 pilins. Scale bar is 1 nm.

Our structural comparison of pilins revealed an additional significant difference between Class 1 and Class 5 pilins. All Class 1 pilins have two cross-linked cysteine residues. This is seen for the known pilin structures, PapA and FimA, and for the homology-modeled pilins, CsnA and FocA. While the precise location of the residues was not preserved in the pri-mary sequences (see Supporting Information Table S1), each disulfide bond cross-links together two intra-subunit *β*-strands, with the approximate disulfide bond location being between cysteines at residues 20 and 60. For example, in PapA (pdb: 5FLU) this S-S bond is between Cys22 and Cys61. In contrast, none of the Class 5 pilins analyzed had any cysteines in their primary sequences, and therefore no S-S linkages could be formed within these pilin subunits.

Although we observed the presence of the short *α*-helix and S-S bond in Class 1 pili, the relevance of these structural features for the response of pili to force was not understood. Therefore, we turned to sMD simulations to investigate the role of specific structural features that affect pilus deformation.

### C. sMD simulations reveal the role of the *α*-helical motif and disulfide bond in Class 1 pilus unwinding

To further investigate the molecular-scale response of pili to tensile force, we carried out sMD simulations of one pilus type from UPEC (P pili) and one from ETEC (CFA/I) based on available cryo-EM structures (Figure 4A, B; (13)(15)). To allow us to focus on the structural changes that occur in region III of unwinding, we used a 3mer system, which eliminates the influence of layer-to-layer interactions that are disrupted in region II (Figure 2). This also minimized the size of the simulated systems, allowing for the use of explicit water solvation and pulling speeds of 5 Å/ns and 1 Å/ns (25, 33–35). We performed three repeats of the sMD simulation at each velocity. As described in more detail in the Methods section, we applied the steering force at a constant velocity along the z-direction, which is aligned with the 3mer central axis (Figure 4C, D). We present data here from one repeat of the slower pulling speed simulations for the P pilus and one for the CFA/I pilus system (1 Å/ns; run 1 in Table S2). Movies from all of the simulations are shown as supplemental movies 1-12, and a full list of simulations and additional information can be found in Table S2 and the Supporting Information.

**Fig. 4.**
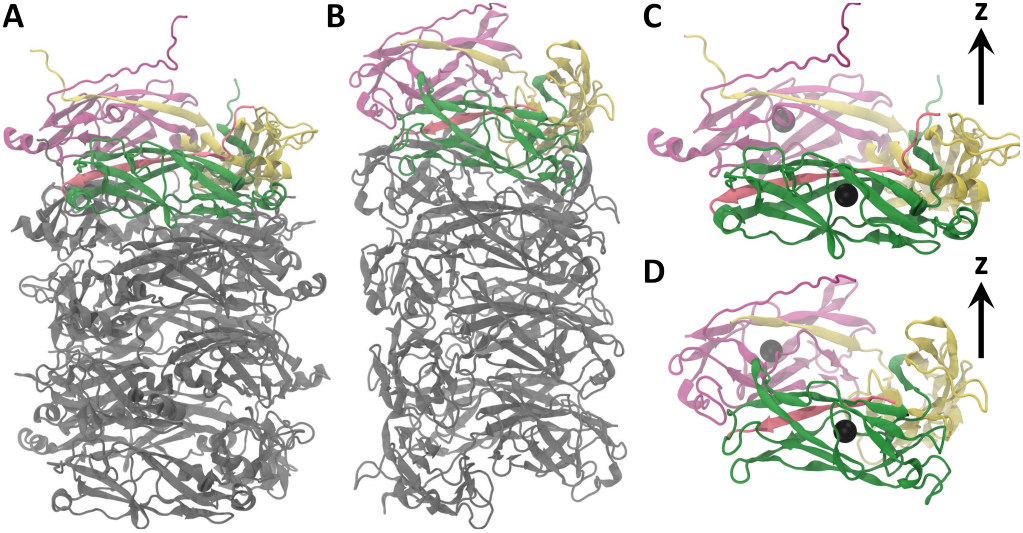
(A) P pilus (5FLU) filament with the simulated subunits colored green, yellow, and pink; also included are the first 20 amino acids of the preceding subunit (red) that make up the inserted *β*-strand of the first (green) subunit. (B) CFA/I pilus (6NRV) filament with the simulated subunits colored green, yellow, pink, and the first 13 amino acids of the preceding subunit (red) that make up the inserted *β*-strand of the first (green) subunit. Remaining subunits in each case are colored in grey and are not included in the simulations. (C) The P pilus 3mer and (D) the CFA/I pilus 3mer, with each panel showing the positions of the centers of mass of the atom selections described in Methods. These centers of mass define the pulling groups in the sMD simulations. The z directional arrow depicts the direction along which pulling forces are applied. Image was made using Visual Molecular Dynamics (VMD) (36).

#### C.1. α-helical unfolding as the source of region III conformational changes in Class 1 pili

During the sMD simulations of the P pilus 3mer we observed that a short segment of amino acids that is in an *α*-helical conformation in the static structure (residues 26-29), undergoes a conformational change and becomes extended near the first observed large force peak (Figure 5A, B, D). While this short helix is a structural element of every subunit in the 3mer of P pili (and in the full filament), we observed that unfolding of this helix only occurred in the middle subunit in our 3mer system. This is consistent with the middle subunit bearing direct strain from pulling, as it is situated between the pulled subunit at the tip of the 3mer and the restrained subunit at the base of the 3mer. Figure 5B shows that for the first approximately 75 Å of 3mer extension the short helix remained relatively unchanged in length. As the peak force was reached the helix extends, unfolding rapidly over approximately 10 Å of 3mer elongation (i.e., from about 85 Å to 95 Å). Once the force again drops (corresponding to a separation event occurring in the 3mer) the helix begins to recover (Figure 5A, B and Figures S1 and S2) as the strain has been removed from the central subunit. We never observe full refolding of this short helical element, as refolding timescales are longer than accessible timescales in sMD simulations. We observed this same trend in the extension of the short helix in the P pilus for every simulation run at both pulling speeds (Figure S2). The observed helix extension was also independent of whether separation of the 3mer occurred between the first and second, or between the second and third subunits, and occurred before a separation of the 3mer (e.g., see Figure S3A). Figure 5D shows several snapshots along the elongation pathway of the P pilus 3mer, highlighting the helical element in purple (see also Supporting Information Movies S1-S6). In contrast to Class 1 pili, ETEC Class 5 pili do not include the sigmoidal shape in region III of unwinding (Figure 2) and also do not contain a corresponding alpha helix in the analogous location. Instead, we observed in CFA/I either a separation event or significant breakdown of tertiary structure (see Supporting Information Movies S7-S12).

**Fig. 5.**
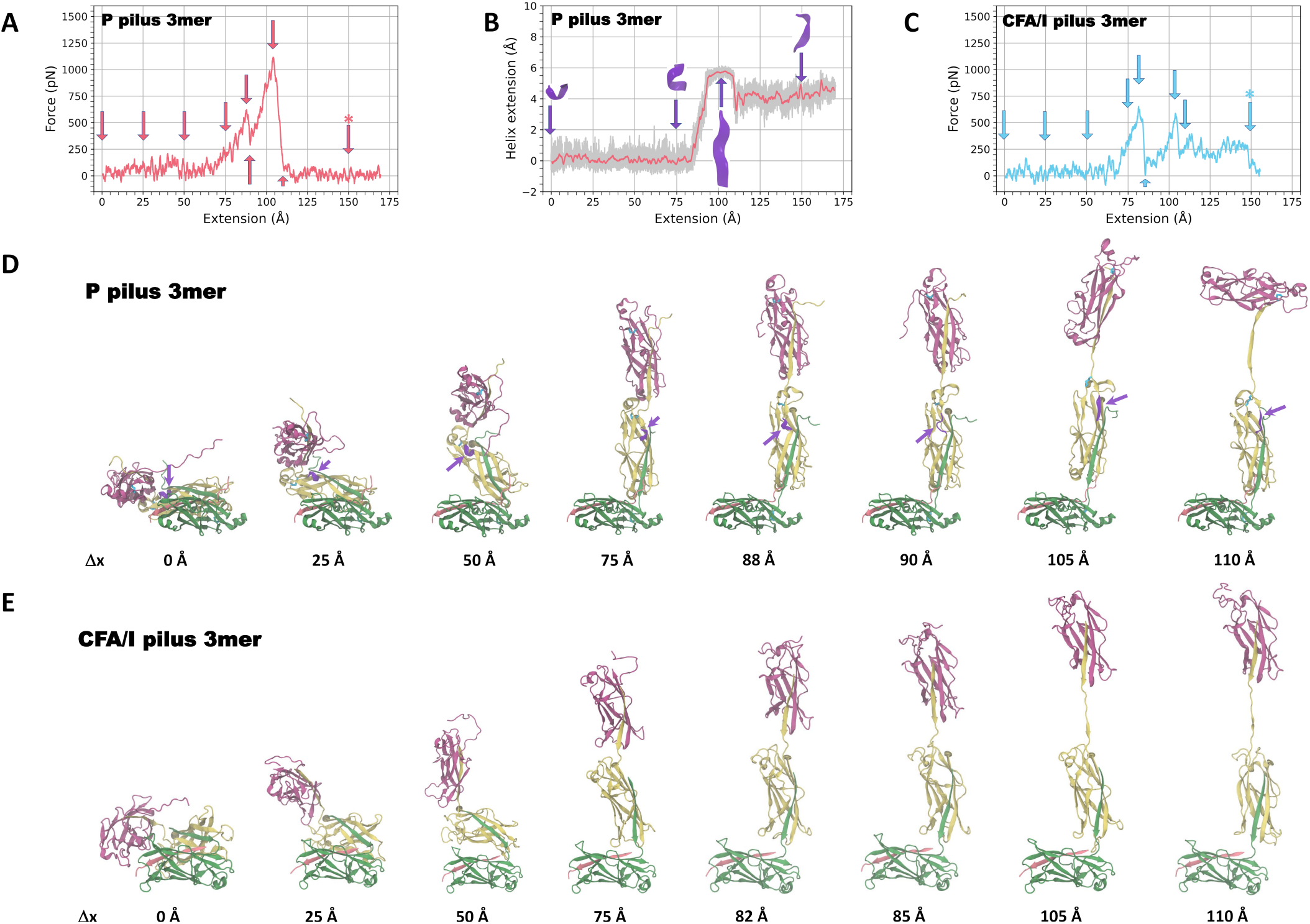
Tensile force applied in the sMD simulations to pili as a function of the 3mer extension. Force is applied along the axial direction, as depicted in Figure 4. (A) For the P pili system (pink), representative data are shown from one simulation at 1 Å/ns pulling speed (run 1 from Table S2). Pink arrows depict the positions at which snapshots of the P pilus are shown in panel D. The image corresponding to the arrow with the asterisk is in Supporting Information Figure S3. The force curve data are a 1 ns running average. (B) Extension of the short *α*-helix in the P pilus as a function of the overall P pilus extension. Several snapshots of the *α*-helical structure (purple) are shown along the trajectory. The pink trace is a smoothed running average (over 1 ns windows) of the raw data (grey). Data are shown from the simulation in panel A. (C) Force as a function of the 3mer extension for the CFA/I pili system (blue); representative data are shown from one simulation at 1 Å/ns pulling speed (run 1 from Table S2). Blue arrows depict positions at which snapshots of the CFA/I pilus are shown in panel E. The image corresponding to the arrow with the asterisk is in Supporting Information Figure S3. (D) Several snapshots from the P pilus 3mer simulation showing the process of elongation of the system under force. The short *α*-helix is shown in purple (marked with purple arrows), and the disulfide bond in each subunit is shown in cyan. (E) Several snapshots from the CFA/I pilus 3mer simulation showing the process of elongation of the system under force.

#### C.2. Simulations suggest that the disulfide bond in Class 1 pili helps to preserve the pilin subunit structure under large forces

Comparing the force-extension profiles of P pili and CFA/I pili from sMD simulations led to several key observations. First, we observed that the peak force obtained in sMD simulations of the P pilus 3mer was consistently higher than the peak force obtained in sMD simulations of the CFA/I pilus 3mer; see Figure 5A, C and Figure S1. Generally, the features of the force-extension profiles for a given system (P pili or CFA/I pili) and pulling speed (5 Å/ns or 1 Å/ns) were highly consistent across each simulation run until the first large force peak (Figure S1). After the first force peak, we observed some subsequent variability in the force-extension profiles between simulation runs. For example, for the P pili system, the green and pink curves at 5 Å/ns and the pink curve at 1 Å/ns showed a slower drop towards 0 pN (Figure S1), corresponding to simulations in which separation of the 3mer occurred between the middle subunit and the bottom subunit. The remaining P pili simulation runs (the blue curve at 5 Å/ns and the blue and green curves at 1 Å/ns) showed a more rapid drop towards 0 pN, and in those simulations separation of the trimer occurred between the middle subunit and the top subunit. In the majority of P pilus 3mer simulations, the tertiary structure of the individual subunits was largely preserved under tensile forces (Figure S3A and Movies S1-S6).

A larger variability between simulation runs was notable in the CFA/I simulations (Movies S7-S12). After the initial elongation that began at an extension of approximately 60-75 Å (similar to P pili), we observed that either the top subunit of the 3mer eventually separated from the system or that there was never a full separation event. In some cases the elongation of the 3mer system was accompanied by substantial disruption of the secondary and tertiary structure of the central subunit. For example in the CFA/I pilus 3mer, unlike P pili at a similar extension length of approximately 105 Å, the central subunit had more extensive disruption of its tertiary structure. At further extension, the tertiary structure of the central subunit was almost fully unraveled (Figure S3B). In the majority of our CFA/I pili simulations, we observed complete separation between two short *β*-strands. This total separation could not occur at the corresponding location in P pilins, where these *β*-strands are restrained by the disulfide bond to remain close together (Figure S4 and S5).

#### C.2. Contribution of the α-helical motif and disulfide bonds to subunit extensibility

To further characterize the effect of the disulfide bond on the change in 3mer structure under force in P pili compared to its absence in CFA/I pili, we also monitored the Val18-Ser39 or Ser10-Ser34 distance in P pili or CFA/I pili, respectively (Figures S6 and S7). We observed that in P pili, where a disulfide bond is present, the maximum change in distance that can occur between Val18-Ser39 after subunits were reoriented and aligned with the pilus axis, was restricted to approximately 16-18 Å in each simulation (Figure S6). However, for the CFA/I pili in which there is no disulfide bond, many simulations resulted in structural failure under high force (e.g., see Figure S3B and S5). Therefore, to compare the extensibility of CFA/I pili to P pili we looked at the two simulations in which the CFA/I pilins did not undergo structural failure. In CFA/I pili we measured the extension of the region Ser10-Ser34 (Figure S7 C,F) to be approximately 7-10 Å. This results in about a 1.6-2.6 fold larger extension of the Val18-Ser39 region of P pili compared to the Ser10-Ser34 region of CFA/I pili from the point where the systems are reoriented and aligned with the pilus axis (taken to be approximately 75 Å of 3mer extension in each case, see Figure 5D,E).

## 2. Discussion and Conclusion

There is a striking similarity in the shapes of the optical tweezers force-extension curves observed in regions I and II of all measured UPEC and ETEC pilus types. We attribute this to similar responses of pilus quaternary and tertiary structures during initial extension. That is, in region I the quaternary structures of pili are stretched, and in region II layer-to-layer interactions are broken, resulting in the sequential unwinding of the helical filament into a linear polymer. It is in region III that differences in unwinding are manifest. In region III, all Class 1 pili, UPEC and ETEC, exhibit an S-shaped extension curve (Figure 2A-C & F) that indicates a rapid stochastic conformational change in the pilus structure after the pilus has been extended to a linear polymer. This conformational change provides a significant increase in length of the pili, amounting to about 25 % of the total extension (Figure 1A).

The source of this conformational change is suggested by structural analysis and sMD simulations of Class 1 and Class 5 phenotypes. Structural analysis reveals an *α*-helix in all Class 1 pili rods that is not present in Class 5 pili. This *α*-helical motif is approximately 20 amino acids from the N-terminal and can unfold under presence of force. Our MD simulations indicate that this unfolding provides flexibility of the subunit, allowing *β*-strands to slip with respect to each other, resulting in as much as a 35 % increase of the PapA subunit’s length at high force. This extension is four-fold in comparison to CfaB subunits. Thus, MD simulation data confirm the additional flexibility observed in region III of P but not in CFA/I pili in optical tweezers force experiments.

Experimental data show that region III of Class 1 pili is fully reversible (3). Consistent with these data, we observe in our sMD simulations of the P pili 3mer that extension of the short *α*-helix in the central subunit begins to shorten spontaneously when force is reduced (Figure 5A, B and Figure S1, S2). This further supports elastic unfolding/refolding of the *α*-helix, and therefore we expect that helix unfolding/refolding is a reversible transition in the structure of the P pilin subunits. This perfectly fits previous force spectroscopy data which show that P pili can be uncoiled and recoiled many times without any sign of fatigue (27).

Under high tensile forces, we see from our sMD study that Class 1 pili most often have a subunit detach from the 3mer (Movies S1-S6). This can occur between the central and top subunit or between the central and bottom subunit in our 3mer systems, and is a result of unzipping of the N-terminal extension *β*-strand that provides the strong, non-covalent interactions between the n and n+1 subunits (see also simulations of unzipping in (37)). This is in contrast to ETEC Class 5 pili, where the observed behavior in our sMD simulations reveals that either a subunit separation event occurs or that the central subunit significantly unravels its secondary and tertiary structure at late times under force (Figure 5 and Figure S3, and Movies S7-S12). Looking comprehensively at the structural and simulation data, this difference in behavior appears to be due to the presence (in Class 1; Figure S4) or absence (in Class 5; Figures S3, S5) of a disulfide bond between *β*-strands within a subunit. The disulfide cross-linking of beta strands in Class 1 pili have a stabilizing effect on the subunit that keeps the subunit more or less intact as the pilus extends to longer lengths, thereby preferring 3mer separation instead of structural failure of the central subunit (Figure S4). These findings agree well with a previous study on the mechanical stability of Type 1 monomers in which the presence of a disulfide bond increased pilin stability significantly (25). In contrast, a subunit that lacks this disulfide bond is less stable. These data support our sMD results for Class 5 pilins that lack cysteine residues (and therefore a disulfide bond), where the *β*-strands tend to pull apart under lower force than Class 1 pilins (Figure S5).

Bacteria that express the Class 1b CS20 pili colonize the intestinal tract and cause diarrheal disease, similar to bacteria expressing CFA/I and CS2. In contrast, the preferred niche of pathogenic bacteria expressing P, Type 1, and F1C pili is the urinary tract. Therefore, it is interesting that the Class 1b ETEC pili CS20 exhibit the conformational change and similar subunit resilience seen in UPEC pili, while requiring lower unwinding forces, more similar to those of Class 5 ETEC pili. This suggests that CS20 pili have adapted to the intestinal tract and that sequence similarity does not correlate directly with environment, as the CS20 major pilin has the lowest homology to the CFA/I major pilin (15 % identity and 36 % similarity), yet both are expressed on ETEC. We, therefore, propose that our findings support the idea of convergent evolution for adhesion pili (38).

Finally, we conclude that despite similar quaternary and tertiary structures, Class 1 and Class 5 pili show differences at the primary (cysteine disulfide bonds) and secondary (*α*-helix) levels that are the basis of significant differences in their biophysical properties. Class 1 pili are more flexible and their subunits tolerate more tensile force, both during unwinding and also after the helical pilus filament has unwound. While the benefit of this extra elasticity for Class 1 ETEC has yet to be explored, we believe this is advantageous for pili expressed on UPEC in the urinary tract, where they encounter high velocity fluid flow. Class 5 ETEC pili, lacking this pilin elasticity, are adapted for the intestinal tract, where they experience lower fluid velocities.

## 3. MATERIALS AND METHODS

### A. Bacterial strains and growth conditions

To avoid possible interference of other bacterial surface organelles, we expressed the UPEC related P pili, Type 1 pili, and F1C using the afimbriated *E. coli* strain HB101, from plasmids pHMG93, pPKL4 and pBSN50, respectively (39–41). We cultured the bacteria on trypticase soy agar at 37 °C. To express ETEC associated CFA/I and CS2 pili we used strains HMG11/pNTP11927 and C91F, respectively (42, 43). Bacteria were grown on CFA agar plates at 37 °C overnight. In addition, CS20 fimbriae were expressed from WS7179A-2/pRA101. Bacteria were grown on Luria-Bertani agar plates with 50 mg ml^-1^ Kanamycin at 37 °C for 24 h (23).

### B. Optical tweezers force measurements

To apply force to pili and measure the corresponding biophysical properties, we used an optical tweezers (OT) setup built around an inverted microscope (Olympus IX71, Olympus, Japan) equipped with a water immersion objective (model: UPlanSApo60XWIR 60X N.A. = 1.2; Olympus, Japan) and a 1920 x 1440 pixel CMOS camera (model: C11440-10C, Hamamatsu) (44). We minimized the amount of noise in the setup and optimized the measured time series using the Allan variance method (45). We used the Power Spectrum method to calibrate the trap by sampling the microspheres position at 131,072 Hz and averaging 32 consecutive data sets acquired for 0.25 s each (46). To extend a pilus, we moved the piezo stage at a constant speed of 100 nm/s for P-pili, F1C, CFA/I, CS2, CS20 and 5 nm/s for Type 1. We sampled the force and position at 100 Hz. We show an illustration of the system in Figure S8A with additional information of the setup in the Supporting Information.

### C. Structural analysis

Structures were taken from the pdb, or for pilins without known structures, the major pilin was homology-modeled using Modeller software (47) and dssp in UCSF ChimeraX (32), after alignment of the genetic sequences using EMBOSS Needle BLOSSUM62 with end gap penalty ‘True’ (48).

### D. Molecular dynamics simulations

We used the *E. coli* P pilus (PDB ID: 5FLU) (13) and the E. coli CFA/I pilus (PDB ID: 6NRV) (15) structures for MD simulations. For each system, we simulated a trimer of pilus subunits extracted from the experimental structures (Figure 4) in which the disulfide bonds were included. All simulations were carried out with the Amber18 software (49) and performed using constant velocity pulling at speeds of 1 Å/ns and 5 Å/ns, with three simulations carried out at each speed (Table S2). The pulling force was applied in the z-direction, extending the pink monomer away from the green monomer in Figures 4 and 5. Additional details are described in the Supporting Information.

## ACKNOWLEDGMENTS

This work was supported to MA by the Swedish Research Council and from the Kempestiftelserna. JLB acknowledges support under NSF grant MCB-1817670. JLB also acknowledges use of the ELSA high performance computing cluster at The College of New Jersey for conducting the simulations reported in this paper. This cluster is funded in part by the NSF under grant numbers OAC-1826915 and OAC-1828163. The authors acknowledge the facilities and technical assistance of the Umeå Core Facility for Electron Microscopy (UCEM).

