## Supporting information for "Impact of an alpha helix and a cysteine-cysteine disulfide bond on the resistance of bacterial adhesion pili to stress"

**This PDF file includes:**
Supplementary text
Sequences
Table S1 and S2
Figs S1 to S8
Supporting Methods text
SI References

**Other supplementary materials for this manuscript include the following:**
Movies S1 to S12

### Supporting Information Text

#### Sequences

All are mature major pilin sequences from UniProtKB.

#### <u>Uropathogenic *E. coli* (UPEC)</u>

>sp|P04127|23-185. **PapA, P pili**
APTIPQGQGKVTFNGTVVDAPCSISQKSADQSIDFGQLSKSFLEAGGVSKPMDLDIELVN
CDITAFKGGNGAKKGTVKLAFTGPIVNGHSDELDTNGGTGTAIVVQGAGKNVVFDGSEGD
ANTLKDGENVLHYTAVVKKSSAVGAAVTEGAFSAVANFNLTYQ

>sp|P04128|24-182. **FimA, Type 1 pili**
AATTVNGGTVHFKGEVVNAACAVDAGSVDQTVQLGQVRTASLAQEGATSSAVGFNIQLND
CDTNVASKAAVAFLGTAIDAGHTNVLALQSSAAGSATNVGVQILDRTGAALTLDGATFSS
ETTLNNGTNTIPFQARYFATGAATPGAANADATFKVQYQ

>tr|Q6KDA6|24-180. **FocA, F1C pili**
AVTTVNGGTVHFKGEVVNAACAVNTNSFDQTVNLGQV
RSERLKVDGAKSNPVGFTIELNDCDSQVSAGAGIVFSGPAVTGKTDVLALQSSAAGSATN
VGVQITDHTGKVVPLDGTASSTFTLTDGTNKIPFQAVYYATGQATAGIANADATFKVQYQ

#### <u>Enteropathogenic *E. coli* (ETEC)</u>

>sp|E3PPC4|24-170. **CfaB, CFA/I pili**
VEKNITVTASVDPVIDLLQADGNALPSAVKLAYSPASKTFESYRVMTQVHTNDATKKVIV
KLADTPQLTDVLNSTVQMPISVSWGGQVLSTTAKEFEAAALGYSASGVNGVSSSQELVIS
AAPKTAGTAPTAGNYSGVVSLVMTLGS

>tr|Q47117|24-170. **CotA, CS2 pili**
AEKNITVTASVDPTIDLMQSDGTALPSAVNIAYLPGEKRFESARINTQVHTNNKTKGIQI
KLTNDNVVMTNLSDPSKTIPLEVSFAGTKLSTAATSITADQLNFGAAGVETVSATKELVI
NAGSTQQTNIVAGNYQGLVSIVLTQEP

>tr|Q8VL73|23-195. **CsnA, CS20 pili**
APAANDSSQATLNFSGRVTSSLCQVKTDDLVKNISLGEVSKSALEATGKSPAQSFQVNLI
NCDSLTDDISYVLADANNNGTTTAYLVPKSGDTAATGVGVFVETSKGTPVNIGSDQKLDV
VANKGNALSEQVIPLRAYIGTQTRAAGAIGTDVTAGTVDATGVLTIRAADATP

**Table S1**

| Pilus | Niche | Class | Major Pilin | UniProtKB ID # | PDB ID | S-S bond | Alpha Helix | Alpha Helix Residue #s |
|---|---|---|---|---|---|---|---|---|
| P pili | UPEC | Class 1 | PapA | P04127 | 5FLU | 22-61 | QKSA | 26-30 |
| Type 1 | UPEC | Class 1 | FimA | P04128 | 6C53, 2JTY | 21-61 | AGSV | 25-28 |
| F1C | UPEC | Class 1 | FocA | Q6KDA6 | | 21-61 | TNSFD | 25-29 |
| CFA/I | ETEC | Class 5 | CfaB | E3PPC4 | 6NRV | none | none | |
| CS2 | ETEC | Class 5 | CotA | Q47117 | | none | none | |
| CS20 | ETEC | Class 1b | CsnA | Q8VL73 | | 23-62 | TDDL | 17-20 |

**Table S2**

| Simulated system | Pulling speed | Run number | Time to simulation end |
|---|---|---|---|
| PapA | 1 Å/ns | 1 | 170 ns |
| PapA | 1 Å/ns | 2 | 170 ns |
| PapA | 1 Å/ns | 3 | 170 ns |
| PapA | 5 Å/ns | 1 | 32 ns |
| PapA | 5 Å/ns | 2 | 34 ns |
| PapA | 5 Å/ns | 3 | 32.8 ns |
| CfaB | 1 Å/ns | 1 | 156 ns |
| CfaB | 1 Å/ns | 2 | 170 ns |
| CfaB | 1 Å/ns | 3 | 160 ns |
| CfaB | 5 Å/ns | 1 | 34 ns |
| CfaB | 5 Å/ns | 2 | 30.8 ns |
| CfaB | 5 Å/ns | 3 | 34 ns |

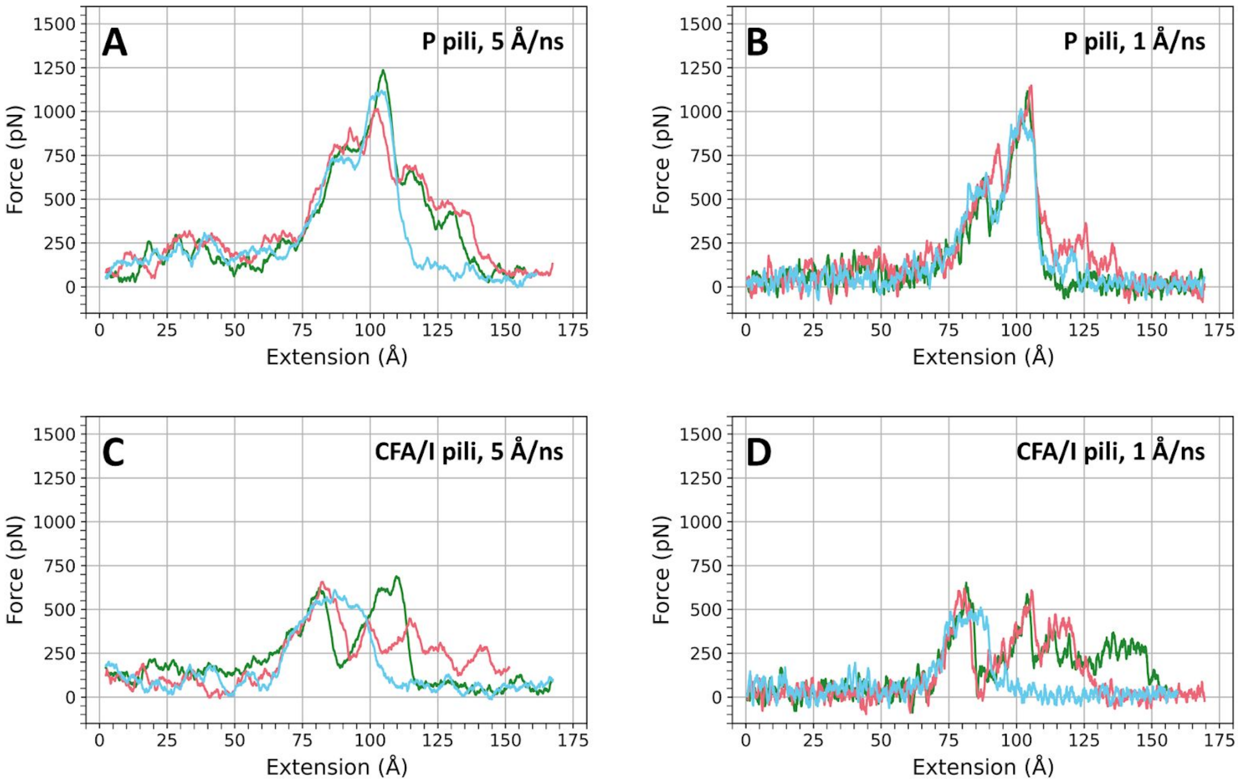


**Figure S1.** Tensile force applied in the sMD simulations to pili as a function of the 3mer extension for the P pili (panels A and B) and CFA/I pili (panels C and D) systems. Panels A/C and B/D are data for the 5 Å/ns and 1 Å/ns pulling speed simulations, respectively. The colors represent the simulation runs (run 1 = green, run 2 = pink, run 3 = blue). The green curve in Panel B and the green curve in Panel D are the same as the pink and blue curves in Figure 5A of the main text, respectively. Force is applied along the axial direction, as depicted in Figure 4 in the main text. The force curves are 1 ns running averages.

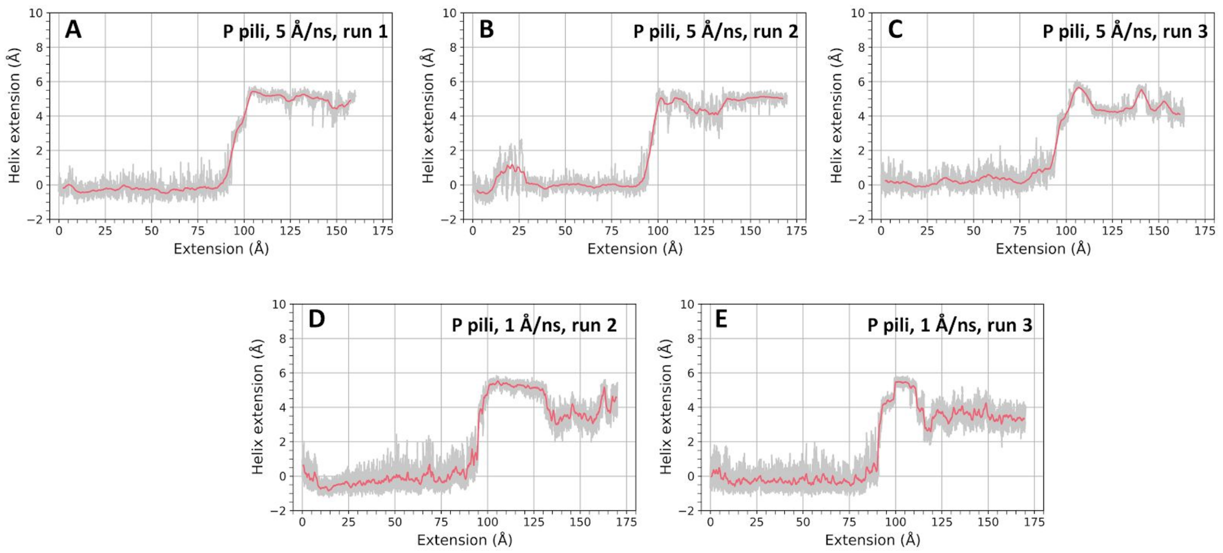


**Figure S2.** Extension of the short α-helix in the P pilus as a function of the overall P pilus extension for (A) 5 Å/ns run 1, (B) 5 Å/ns run 2, (C) 5 Å/ns run 3, (D) 1 Å/ns run 2, and (E) 1 Å/ns run 3. The data for 1 Å/ns run 1 is provided in the main manuscript, Figure 5B. The pink trace is a smoothed running average (over 1 ns windows) of the raw data (grey).

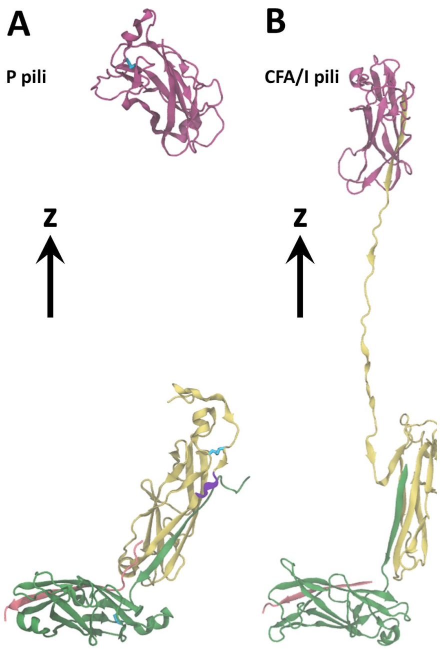


**Figure S3.** Snapshot from 150 Å extension from 1 Å/ns pulling trajectory of the (A) P pili 3mer (run 1) and the (B) CFA/I pili 3mer (run 1). In this run the P pili terminal monomer separates from the 3mer, while the CFA/I terminal monomer remains attached by the central monomer and is dramatically distorted/unfolded. The central monomer's dramatic disruption of its secondary and tertiary structure is attributed to the absence of a disulfide bond in the CFA/I system compared to the P pili system. In the P pili 3mer the disulfide bonds are shown in cyan licorice representation. The z directional arrow depicts the direction along which pulling forces are applied in the simulations.

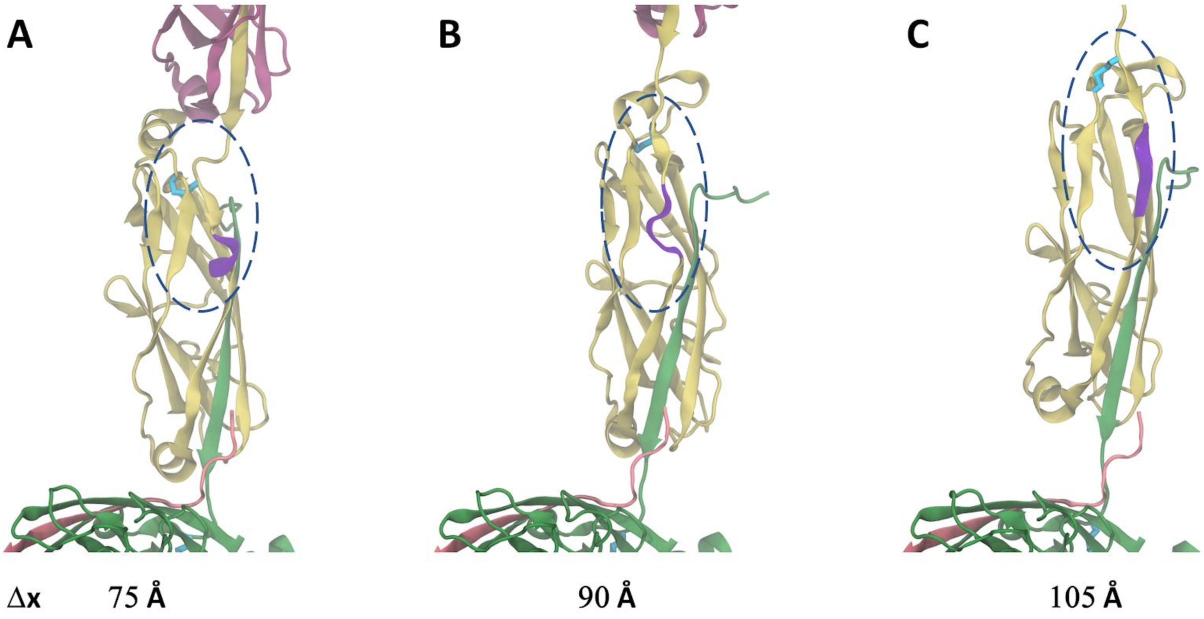


**Figure S4.** Snapshots from the P pili 3mer 1 Å/ns pulling trajectory (run 1) showing extension lengths of (A) 75 Å, (B) 90 Å, and (C) 105 Å. The disulfide bond is shown in cyan, the short helix is shown in purple, and the blue-dashed region highlights the beta strands of interest. The beta strands slip away from one another under force as seen in the 105 Å snapshot, however the disulfide bond helps to preserve the overall structure of that connected region.

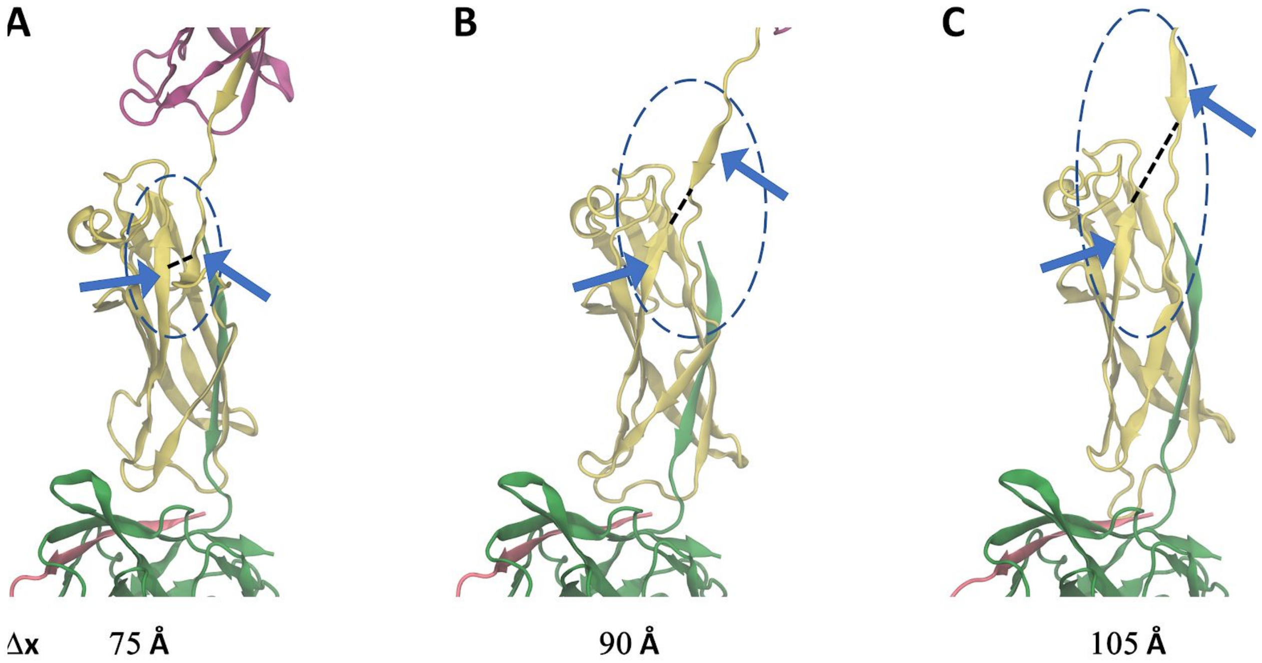


**Figure S5.** Snapshots from the CFA/I pili 3mer 1 Å/ns pulling trajectory (run 1) showing extension lengths of (A) 75 Å, (B) 90 Å, and (C) 105 Å. The blue-dashed region highlights the beta strands of interest. Blue arrows point to the beta strands. The black dashed line is used to emphasize the separation between these structural elements of the CFA/pili central subunit. The beta strands slip away from one another under force as seen in the 90 Å and 105 Å snapshots, and in the absence of a disulfide bond they are free to continue to separate as the system is extended, disrupting the overall structure of the central monomer.

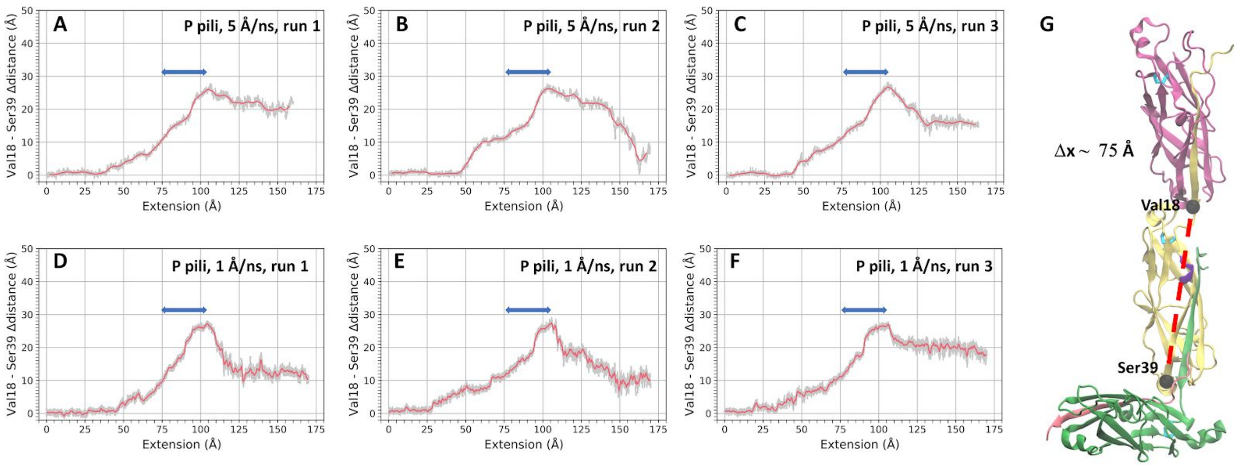


**Figure S6.** (A) - (F) show the change in the distance (relative to the initial distance) between the alpha carbons of amino acids Val18 and Ser39 in the central subunit of the P pili 3mer (see the dark beads in (G)) as a function of the 3mer extension. This is the same 3mer extension used for the abscissa in Figure 5 of the main manuscript and in Figures S1 and S2 above. This distance can be used to estimate approximately how much extension occurs within a single pilin subunit as the entire 3mer is extended. The image in G is a snapshot from the 1 Å/ns run 1 simulation with an Extension value of ~ 75 Å. At ~ 75 Å of 3mer extension the 3mer is at approximately a 90 degree angle as pictured in G. The blue arrow in each image goes from this point at ~ 75 Å until we start to see a decrease in the Val18-Ser39 distance change. Over the length of the blue arrow there is an additional increase in the Val18-Ser39 distance change of ~ 16-18 Å. The pink trace is a smoothed running average (over 1 ns windows) of the raw data (grey).

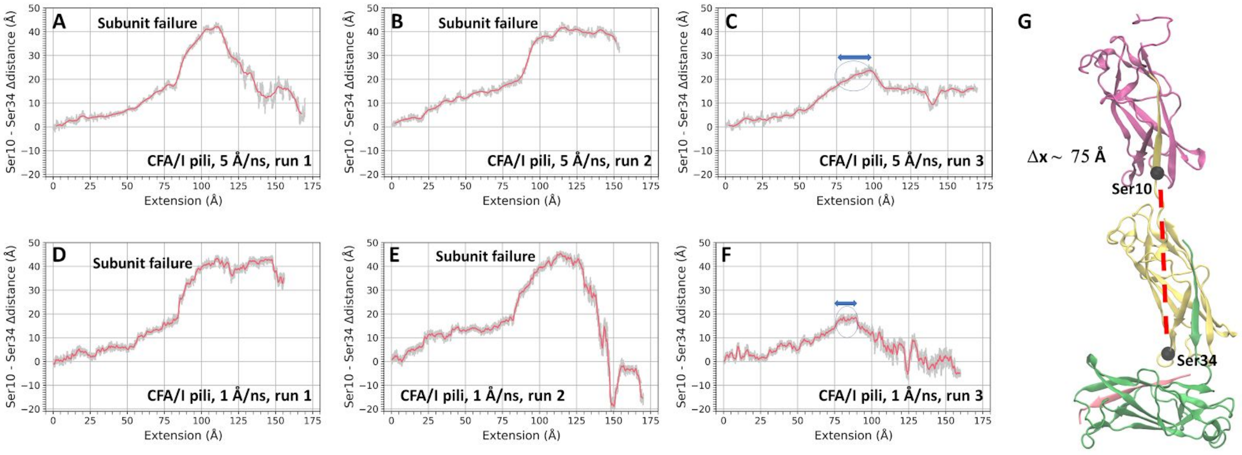


**Figure S7.** (A) - (F) show the change in the distance (relative to the initial distance) between the alpha carbons of amino acids Ser10 and Ser34 in the central subunit of the CFA/I 3mer (see the dark beads in (G)) as a function of the 3mer extension. This is the same 3mer extension used for the abscissa in Figure 5 of the main manuscript and in Figures S1 and S2 above. This distance can be used to estimate approximately how much extension occurs within a single pilin subunit as the entire 3mer is extended. The image in G is a snapshot from the 1 Å/ns run 1 simulation with an Extension value of ~ 75 Å. At ~ 75 Å of 3mer extension the 3mer is at approximately a 90 degree angle as pictured in G. We note that the additional change in the Ser10-Ser34 distance is highly variable in the different simulations depending on how the trajectory progresses (e.g., see the corresponding Supplemental Movies). In simulations in which subunit failure occurs (meaning that the beta strands in Figure S5 separate allowing for significant unraveling of the CFA/I middle monomer, e.g., panels A, B, D, E) a very large extension of greater than 40 Å is seen due to the failure. For two of the CFA/I simulations, failure does not occur, corresponding to panels C and F. The blue arrow in panels C and F image goes from ~ 75 Å (the point at which the 3mer is oriented at approximately 90 degrees) until we start to see a decrease in the Ser10-Ser34 distance change. Over the length of the blue arrow there is an additional increase in the Val10-Ser34 distance change of ~ 7-10 Å. This is notably smaller than the additional distance change observed for the Val18-Ser39 distance in Figure S6. The pink trace is a smoothed running average (over 1 ns windows) of the raw data (grey).

### Methods

#### Optical tweezers (OT) instrumentation and sample preparation

The OT stands in a temperature controlled room with computers and controllers isolated from the room to reduce noise and vibrations. We use a continuous wave DPSS laser (Cobolt Rumba, Cobolt AB, Solna, Sweden) operating at 1064 nm for trapping a single bacterium or microsphere. A HeNe probe laser operating at 632.8 nm, (117A, Spectra Physics, Santa Clara, CA) is merged with the trapping laser using a dichroic mirror (DMSP650, Thorlabs, Newton, NJ). The light from the probe laser is refracted by the trapped object and collected by the condenser and thereafter imaged onto a 2D position-sensitive detector (PSD, 2L10YAG SU65 SPC02, Sitek Electro Optics, Sweden). The PSD converts the incoming light to a photocurrent and thereafter to a voltage that is sent to a programmable low pass filter (SR640, Stanford research systems), later collected by a computer and processed with an in-house LabVIEW program.

To prepare a sample we suspended bacteria in 1xPBS to a concentration (1:1000 of OD600 = 1) suitable for single cell analysis using optical tweezers. Surfactant-free 2.0 μm amidine polystyrene microspheres (product no. 3-2600, Invitrogen, Carlsbad, CA) were similarly suspended in Milli-Q water; these microspheres were trapped and used as force probes. To mount bacteria and reduce the influence of surface interactions we prepared a 1:500 suspension of 9.5 μm carboxylate-modified latex microspheres (product no.2-10000, Interfacial Dynamics, Portland, OR) in Milli-Q. We dropped ten μL of the microsphere-water suspension onto 24 x 60 mm coverslips (no.1, Knittel Glass, Braunschweig, Germany) and placed these in an oven for 60 min at 60° C to immobilize the microspheres to the surface. To firmly adhere bacteria to the microspheres, we added a solution of 20 μL of 0.01 % poly-L-lysine (catalog no. P4832, Sigma-Aldrich, St. Louis, MO) to the coverslips, which, after 45 min incubation at 37° C, were stored until use.

To make a flow chamber, we added a ring of vacuum grease (Dow Corning, Midland, MI) around the area containing the poly-L-lysine-coated microspheres on one of the coverslips. Carefully, we dropped a 3-mL suspension of bacteria and a 3-μL suspension of the probe microspheres onto the area and sealed the flow chamber by placing a 20 x 20 mm coverslip (no.1, Knittel Glass) on top. Thereafter, we mounted the sample in a sample holder that was fixed to a piezo-stage (Physik Instrument, P-561.3CD stage) in the OT instrumentation. To get reliable OT calibration parameter values we measured the temperature using a thermistor mounted on the nosecone of the objective, 24.0° C ± 0.1° C. The viscosity was assumed to only vary with temperature and thus set to 0.932 mPas ± 0.002 mPas.

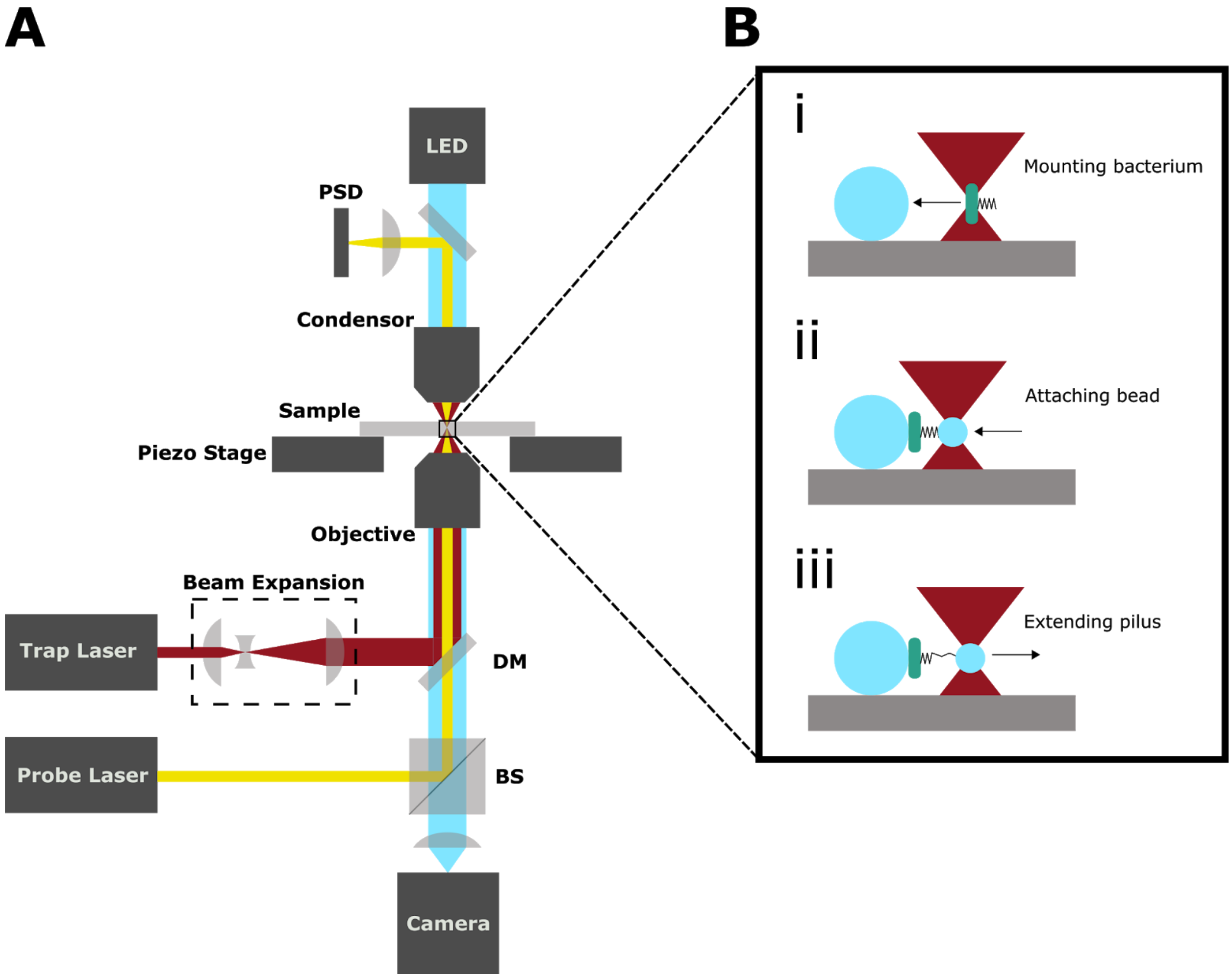


**Figure S8.** Schematic of the optical tweezer setup used to extend pili. (A) We merge the trap and probe laser beams using a beam splitter cube (BS) and a dichroic mirror (DM). The two laser beams are thereafter focused by the objective into the sample. To sample the position of a trapped microsphere we use a weak probe laser beam (μW) that is scattered by the sphere and collected by the condenser. The beam illuminates a PSD that converts the incoming light to a photocurrent, which in turn is converted to a voltage signal and acquired by a computer. (B) An illustration of a force-extension measurement. i) We trap a single bacterium and mount it onto a large microsphere that is immobilized to the cover slip. ii) We trap a small microsphere that we attach to a pilus. iii) We separate the bacterium and trapped microsphere to apply a tensile force to the pilus. This allows us to extend and measure the resistance to extension (force) on individual pili.

**Detailed molecular dynamics simulation methods**

All systems were prepared for simulation using the tLeap program included with AmberTools18 (1). The 3mer systems were already aligned along the z-direction in the initial structure file, and this system orientation was preserved in preparation for simulation. The FF14SB force field was used for protein parameters (2), and the TIP3P force field was used for water (3). The Joung and Cheatham parameters were used for monovalent counterions (4). Both 3mer systems were solvated in a water box with a 12 Å buffer to the edge of the periodic cell in the x and y directions (orthogonal to the 3mer central filament axis), and with a 90 Å buffer along the z-direction in order to allow for sufficient water buffer for the extension of the 3mer system along the filament axis in sMD simulations. To neutralize the overall charge of the solvated systems, 15 Na+ counterions were added to the P pilus system and 7 Na+ counterions were added to the CFA/I pilus system.

To prepare each system for sMD simulations we used the following approach. First, we energy minimized the system using 3000 steps of steepest descent and 2000 steps of conjugate gradient. During minimization, the Cα atoms were restrained using a force constant of 10.0 kcal $mol^{-1}$ $Å^{-2}$. After minimization, the systems were heated in two separate stages. The initial stage of heating was performed using the NVT ensemble, and the system temperature was increased from 0 K to 100 K over 20 ps and then held at 100 K for an additional 30 ps. The second stage of heating was carried out in the NPT ensemble, during which the system temperature was increased from 100 K to 300 K over 20 ps and then held at 300 K for an additional 80 ps. Restraints of 10.0 kcal $mol^{-1}$ $Å^{-2}$ were applied to the Cα atoms during both stages of heating. After heating, systems were equilibrated while using Cα restraints. Equilibration was performed over six stages and carried out in the NPT ensemble at a temperature of 300 K. Each of the first five stages of equilibration were 200 ps each during which restraints on the Cα atoms were reduced from 10.0 kcal $mol^{-1}$ $Å^{-2}$ to 0.1 kcal $mol^{-1}$ $Å^{-2}$ (restraints were reduced as 10.0, 5.0, 2.5, 0.5, and 0.1 kcal $mol^{-1}$ $Å^{-2}$. The final sixth stage of equilibration was 5 ns long and and Cα restraints were placed on the red beta strand, green subunit (except for residues 1-20 for 5FLU and 1-13 for 6NRV), and the pink subunit with a strength of 0.1 kcal $mol^{-1}$ $Å^{-2}$. Three frames were extracted from the end of the last stage of equilibration, separated by 400 ps from one another, to use as the initial structures for the subsequent sMD simulations (5). Specifically, each of the three sMD simulations at each speed started from one of the three final frames from equilibration.

The sMD simulations were carried out using the jar = 1 (6) option in Amber18 and were conducted in the NPT ensemble with a temperature of 300 K. The collective variable (CV) used to describe the fixed and pulled atom groups for the sMD simulations is the z-distance between the Cα atoms of the green subunit and red beta strand (except for residues 1-20 for 5FLU and 1-13 for 6NRV) and the Cα atoms of the pink subunit (except for residues 1-15 for 5FLU and 1-13 for 6NRV). Application of the sMD force along the z-direction was achieved using the fxyz option in Amber18. During the sMD simulations, the green subunit and red beta strand Cα atoms were restrained using a 0.5 kcal $mol^{-1}$ $Å^{-2}$ restraint in each sMD simulation as the system was extended along the CV in order to prevent overall rotations and translations of the system (mimicking the 3mer base being adhered to a surface as the top end is pulled). Each system was extended by approximately 170 Å along the CV which generally resulted in elongation and finally separation of the 3mer subunits from one another. Simulations were ended if the top pilin subunit approached too closely to the bottom pilin subunit in the periodic image of the main simulation cell. The stiffness of the pulling spring was set to 10.0 kcal $mol^{-1}$ $Å^{-2}$. Force versus extension data was saved every 2 ps. For the heating, equilibration, and pulling phases of simulation a 2 fs integration timestep was used, which was made possible by applying the SHAKE algorithm to hydrogen bonds (7). Temperature was regulated using a Langevin thermostat with a collision frequency of 1 $ps^{-1}$ and in NPT simulations constant pressure of 1 atm was maintained using the Monte Carlo barostat (8). A real space interaction cutoff of 8 Å was employed in all stages of the simulations, and the particle mesh Ewald method was used for long range electrostatics(9).

**Supporting Movies**

**Supporting Movie S1.** Movie rendered using VMD of **P pilus 3mer** simulation run 1 **v = 5 Å/ns**. Coloring scheme is the same as in Figure 5 of the main manuscript.

**Supporting Movie S2.** Movie rendered using VMD of **P pilus 3mer** simulation run 2 **v = 5 Å/ns**. Coloring scheme is the same as in Figure 5 of the main manuscript.

**Supporting Movie S3.** Movie rendered using VMD of **P pilus 3mer** simulation run 3 **v = 5 Å/ns**. Coloring scheme is the same as in Figure 5 of the main manuscript.

**Supporting Movie S4.** Movie rendered using VMD of **P pilus 3mer** simulation run 1 **v = 1 Å/ns**. Coloring scheme is the same as in Figure 5 of the main manuscript.

**Supporting Movie S5.** Movie rendered using VMD of **P pilus 3mer** simulation run 2 **v = 1 Å/ns**. Coloring scheme is the same as in Figure 5 of the main manuscript.

**Supporting Movie S6.** Movie rendered using VMD of **P pilus 3mer** simulation run 3 **v = 1 Å/ns**. Coloring scheme is the same as in Figure 5 of the main manuscript.

**Supporting Movie S7.** Movie rendered using VMD of **CFA/I pilus 3mer** simulation run 1 **v = 5 Å/ns**. Coloring scheme is the same as in Figure 5 of the main manuscript.

**Supporting Movie S8.** Movie rendered using VMD of **CFA/I pilus 3mer** simulation run 2 v = **5 Å/ns**. Coloring scheme is the same as in Figure 5 of the main manuscript.

**Supporting Movie S9.** Movie rendered using VMD of **CFA/I pilus 3mer** simulation run 3 **v = 5 Å/ns**. Coloring scheme is the same as in Figure 5 of the main manuscript.

**Supporting Movie S10.** Movie rendered using VMD of **CFA/I pilus 3mer** simulation run 1 **v = 1 Å/ns**. Coloring scheme is the same as in Figure 5 of the main manuscript.

**Supporting Movie S11.** Movie rendered using VMD of **CFA/I pilus 3mer** simulation run 2 **v = 1 Å/ns**. Coloring scheme is the same as in Figure 5 of the main manuscript.

**Supporting Movie S12.** Movie rendered using VMD of **CFA/I pilus 3mer** simulation run 3 **v = 1 Å/ns**. Coloring scheme is the same as in Figure 5 of the main manuscript.
